# Cochlear activity in silent cue-target intervals shows a theta-rhythmic pattern and is correlated to attentional alpha modulations

**DOI:** 10.1101/653311

**Authors:** Moritz Herbert Albrecht Köhler, Gianpaolo Demarchi, Nathan Weisz

## Abstract

A long-standing debate concerns where in the processing hierarchy of the central nervous system (CNS) selective attention takes effect. In the auditory system cochlear processes can be influenced via direct and mediated (by the inferior colliculus) projections from the auditory cortex to the superior olivary complex (SOC). Studies illustrating attentional modulations of cochlear responses have so far been limited to sound-evoked responses. The aim of the present study is to investigate intermodal (audiovisual) selective attention in humans simultaneously at the cortical and cochlear level during a stimulus-free cue-target period. We found that cochlear activity in the silent cue-target periods was modulated by a theta-rhythmic pattern (∼6 Hz). While this pattern was present independently of attentional focus, cochlear theta activity was clearly enhanced when attending to the upcoming auditory input. On a cortical level, classical posterior alpha and beta power enhancements were found during auditory selective attention. Interestingly, participants with a stronger release of inhibition in auditory brain regions show a stronger attentional modulation of cochlear theta activity. These results hint at a putative theta-rhythmic sampling of auditory input at the cochlear level. Furthermore, our results point to an interindividual variable engagement of efferent pathways in an attentional context that are linked to processes within and beyond processes in auditory cortical regions.

## Introduction

Cognitive processing of sensory stimuli is capacity limited. Hence, attentional processes are required to prioritize cognitive resources on task- or context-relevant stimuli. On a neural level, responses to attended stimuli are enhanced, while responses to unattended and distracting stimuli are diminished (Couperus & Mangun, 2010; Fritz et al., 2007). These effects have been mainly established on a cortical level (Frey et al., 2015; Shrem & Deouell, 2017); however, it is less clear to what extent selective attention modulates subcortical activity (Guinan, 2018). For the auditory system, this dispute extends down to the level of the cochlea (Beim et al., 2018; Giard et al., 1994; Lopez-Poveda, 2018).

Indeed cochlear processes can be modulated via direct and mediated (by the inferior colliculus) projections from the auditory cortex to the superior olivary complex (SOC). The SOC finally innervates the outer hair cells (OHC) that are essential for cochlear amplification and fine tuning of the basilar membrane (Delano & Elgoyhen, 2016). The architecture of the efferent auditory system would – in principle – enable the auditory cortex to modulate cochlear processes (Terreros & Delano, 2015).

An increasing number of studies support this notion by measuring otoacoustic emissions (OAE; Smith, Aouad, & Keil, 2012; Walsh, Pasanen, & McFadden, 2015; Wittekindt, Kaiser, & Abel, 2014) or cochlear microphonics (Delano et al., 2007). However, the described effects are restricted to sound-evoked responses, are small, and sometimes contradictory (Francis et al., 2018; Meric & Collet, 1992). Furthermore, the attention research on cortical and cochlear processes has been conducted largely independently (see Wittekindt et al. (2014), Dragicevic et al. (2019), or Riecke et al. (2020) for exceptions). In summary, it remains unclear whether and how attention modulates cochlear processes during silent periods and how these peripheral processes are linked to cortical processes.

We applied an established intermodal (audiovisual) selective attention task and simultaneously measured activity from different levels of the auditory system, to advance our knowledge in this area. To stay as close as possible to previous magnetoencephalography and electroencephalography (M/EEG) works in this domain (Foxe et al., 1998; Frey et al., 2014), we decided to record sounds within the ear canal during silent cue-target periods. This “ongoing otoacoustic activity” (OOA) allows for an unbiased measurement of cochlear modulations by cortical attention processes, since undesired sound-evoked cochlear changes are circumvented (Guinan et al., 2003).

Given that attentional modulations of cortical oscillations are mostly found at low frequencies (< 30 Hz), we decided to use a similar analysis approach for the OOA-signal as Dragicevic et al. (2019), an approach that allows us to investigate oscillatory cochlear activity at the same frequencies as cortical activity occurs. Further, genuine periodic components (peaks) of the OOA-signal were computed for the OOA (Haller et al., 2018). Replicating an established finding from several previous studies (Fu et al., 2001; Klimesch, 2012; Wittekindt et al., 2014), we show strong attentional modulation of visual cortical alpha activity. More importantly, we illustrate a rhythmic modulation of cochlear activity in the theta frequency range. While this theta activity was generally present independently of attentional focus, it was strongly amplified when attending to the auditory modality. Interestingly, this attentional amplification of cochlear activity is inversely correlated with attentional alpha and theta effects at the cortical level across participants.

## Methods

### Participants

34 healthy volunteers (23 females, age range: 18-35 years) participated in this study. One participant was excluded from analyses because his right ear was occluded by cerumen. As recording otoacoustic activity inside an MEG system is challenging, further participants were also excluded from the final analysis (see below in the results section for details). One participant was excluded because the left acoustic meatus was too small to fit the foam ear tip without causing pain. One participant was excluded because the recordings from the left ear showed excessive periods of saturation. Another four participants were excluded because the number of artifact contaminated MEG trials exceeded two standard deviations of the mean. The remaining 27 volunteers (18 female, mean age: 22.96 years, age range: 18-35 years) were used for analyses. Four participants were left handed. None of the participants reported any known hearing deficit and any visual impairment was corrected to normal with MEG-compatible glasses. All subjects were informed about the experimental procedure and the purpose of the study and gave written informed consent. As compensation subjects received either €10 per hour or credit for their psychology studies. This study was approved by the Ethics Committee of the University of Salzburg.

### Stimuli and Procedure

Our focus in this study was to investigate intermodal selective attention by simultaneously measuring cochlear (OOA) and neuronal processes (MEG). Studies investigating attentional modulations of OAEs in the past often used a block design (Froehlich et al., 1993; J. L. Puel et al., 1988; Smith et al., 2012). As this procedure is criticized for not achieving highly controlled attentional conditions (Carrasco et al., 2004; Ward, 1997; Wittekindt et al., 2014), we decided to use an adapted version of the trial-wise cueing paradigm introduced by Wittekindt et al. (2014).

Measurements took place in a magnetically shielded room (AK3B, Vacuumschmelze, Hanau, Germany), in which subjects sat quietly inside the MEG system (TRIUX, MEGIN-Elekta Oy, Helsinki, Finland). Participants performed five blocks consisting of 80 trials (40 Attend Auditory and 40 Attend Visual) in a pseudo-randomized order. **Figure 1** schematically illustrates the course of a trial. Each trial started with a visually presented cue (1 s duration) instructing the subject to either attend the auditory or the visual modality. The letter “A” indicated the Attend Auditory condition and the letter “V” the Attend Visual condition. During the following silent cue-target period (2 s duration) a fixation dot was presented and the participants had to shift their attention selectively to the indicated modality. To eliminate any effects of divided attention and to reach maximum focus on the cued modality, the cue was 100% informative (Wittekindt et al., 2014). The target stimulus in the visual modality was a low-contrast Gabor patch (diameter: ca. 2 degrees of visual angle) that was displayed in the center of a rear projection screen placed inside the shielded room (distance to the subject: 1.1 m) and oriented 45 degrees to the right or left. The target stimulus in the auditory modality was a pure tone of either 1131 Hz or 1987 Hz, which was presented via ear inserts. The sound volume was individually adjusted to be at a comfortable level. Visual and auditory stimuli were simultaneously presented for 100 ms. For the auditory stimuli, we employed two 5 ms linear fade in/out windows. Depending on the preceding cue, the task was to detect the orientation of the Gabor patch (Attend Visual, left or right 45° tilt) or the pitch level of the tone (Attend Auditory, high pitch (1987 Hz) or low pitch (1131 Hz)). Afterwards, a response screen showed indicators for choosing either the pitch level of the tone or the orientation of the Gabor patch. Participants were instructed to wait until the response screen was presented (0.5 s post-target), and then reply as soon as they were ready by pressing the corresponding button with their left or right thumb, within 2 s after the appearance of the response screen. The inter-trial intervals were jittered uniformly between 1 and 2 s. Acoustic and visual stimuli were generated by the Psychophysics Toolbox Version 3 (Brainard, 1997; Pelli, 1997) using custom-written MATLAB scripts (Version 9.1; The MathWorks).

**Figure 1.**
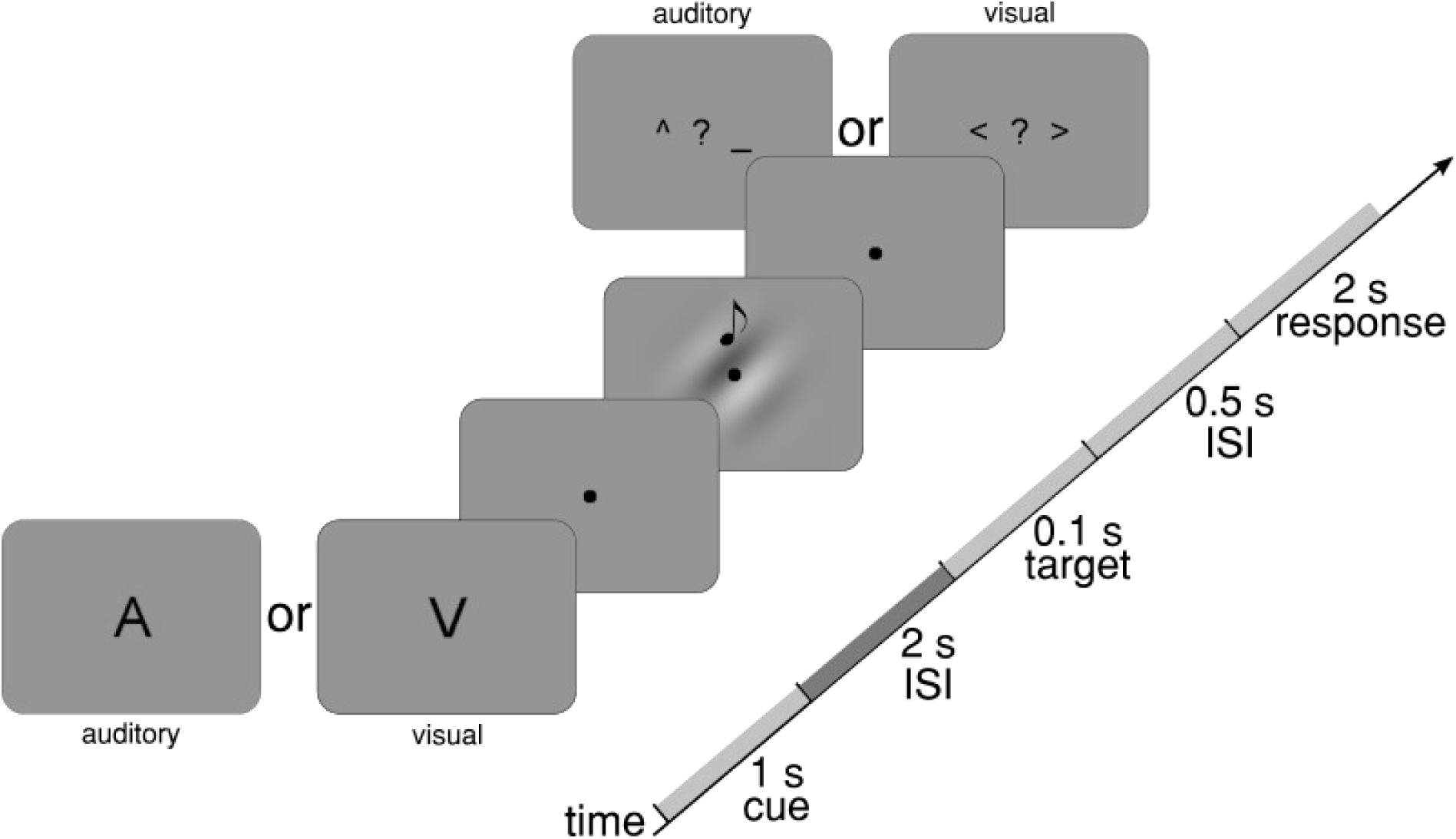
Schematic illustration of the task. Each trial started with a 100% informative visual cue telling the subject to either attend the auditory (“A”) or the visual modality (“V”). After an ISI of 2 s a left or right oriented Gabor patch and a low-frequency (1131 Hz) or high-frequency (1987 Hz) pure tone were simultaneously presented. After another ISI of 0.5 s a response screen depending on the cued modality appeared for 2 s. The intertrial interval was uniformly jittered between 1–2 s.

### Recording of Cochlear and Cortical Activity

In order to measure otoacoustic activity, a probe consisting of a sensitive microphone and two loudspeakers (ER-10C microphone/preamplifier system, Etymotic Research, Elk Grove Village, US) was fitted into the subject’s right and left ear canal with a foam ear tip. Otoacoustic activity was recorded from both ears concurrently. The microphone signal was fed into the EEG amplifier of the MEG system, with an amplitude gain of +55 dB (600x). The sampling rate of the entire MEG and EEG system was set to 10 kHz. The ER-10C received its input via two BNC cables coming from a sound preamplifier (SOUNDPixx, VPixx Technologies, Saint-Bruno, Canada). The SPL for the loudspeakers was balanced to the left and right side by subjective feedback for each participant.

Neuromagnetic brain activity was recorded with 306 channels (TRIUX MEG, see above). Two bipolar electrodes were mounted above and below the left eye, one was mounted on the left side of the left eye and another on the right side of the right eye to monitor eye blinks and eye movements (H/VEOG). Further, two electrodes were mounted on the bottom left rib and the right collarbone to record electrocardiography (ECG). A reference electrode was placed on the left trapezius muscle, and the ground electrode on the right supinator. Prior to the experiment, individual head shapes were acquired for each participant including relevant anatomical landmarks (nasion and preauricular points) and about 300 digitized points on the scalp with a 3D digitizer (Polhemus FASTRAK, Colchester, US). Head positions of the subjects in the helmet were estimated at the beginning of each block injecting a small current into five (HPI, head position indicator) coils. Again, the overall (MEG+EEG) sampling rate was set to 10 kHz, with a hardware high-pass filter of 0.1 Hz, and an anti-alias low-pass filter with the cutoff frequency set to 3330 Hz.

### Signal Processing

OOA was preprocessed by high-pass filtering at 500 Hz (6^th^ order Butterworth IIR), extracting epochs of 3 s duration after cue presentation and manually rejecting trials containing periods of signal saturation or atypical high background noise, for example, caused by moving, swallowing, or coughing (average number of rejected trials per participant: 87.15; range across participants: 1-185). As the frequencies of the acoustic targets were between 1131 Hz and 1987 Hz and otoacoustic activity is strongest in the range from 1000-2000 Hz (Puria, 2003), we expected amplitude modulations of the OOA in this range. The cue-target period was defined as the period in which intermodal attention processes occur (Wittekindt et al., 2014). In a next step, trials were split into two conditions (Attend Auditory and Attend Visual), averaged over 1.7 s of the cue-target period, and bandpass filtered in 10 Hz steps from 1000-2000 Hz (bandpass window +/- 30 Hz). This resulted in 201 bandpass windows for each participant, which represent the binned cochlear frequency response between 1000 and 2000 Hz. To be able to further study any relationship between cochlear activity and brain oscillations (see Results section), we extracted the envelope of the cochlear signal for each of the previous bandpass windows via a Hilbert transform, thus obtaining a signal with a frequency range that is routinely used in electrophysiological evaluations of cognitive tasks. Next, power spectral density (PSD) from 1-30 Hz was calculated for each condition and each Hilbert transformed bandpass window (“mtmfft” fieldtrip implementation with a Hann window). Finally, the bandpass windows were concatenated for each condition resulting in a representation of the amplitude modulation from 1-30 Hz at cochlear response frequencies from 1000-2000 Hz.

The MEG signal was first preprocessed by manually rejecting all bad sensors (average number of rejected sensors per participant: 38.89; range across participants: 13-73), high-pass filtering at 1 Hz (6^th^ order Butterworth IIR), extracting epochs of 3 s duration after cue presentation and down-sampling to 1 kHz. The excessive amount of rejected sensors is caused by magnetic artifacts of the microphone probes, which leads to a saturation of several mostly temporal sensors. The detected bad trials in the OOA data were used to reject the same trials in the MEG data. In a next step trials were again split into two conditions (Attend Auditory and Attend Visual). For source level analysis, a standard anatomical magnetic resonance imaging (MRI) template provided by the Statistical Parametric Mapping toolbox (Version 12; Friston, Penny, Ashburner, Kiebel, & Nichols, 2006) was morphed to the individual head shape of each participant using non-linear-transformation. Sensor space trials were projected into source space using linearly constrained minimum variance (LCMV) beamformer filters (Van Veen et al., 1997). The aligned brain volumes were also used to create single-shell head models and compute the leadfield matrices (Nolte, 2003). For the template grid we chose a resolution of 1 cm in MNI space. PSD in 1 Hz steps in a frequency range of 1-30 Hz averaged over 1.7 s of the cue-target period was calculated for each condition by a FFT (Hann window). The preprocessing of the OOA and MEG data were conducted using the open-source FieldTrip toolbox for EEG/MEG data (Oostenveld et al., 2011) and custom-written MATLAB scripts (Version 9.1; The MathWorks).

### Statistical Analysis

As a first analysis step, we investigated if rhythmic modulations of cochlear activity are present. The python (Version 3.7.1) toolbox FOOOF (Haller et al., 2018) was used to parameterize the power spectra of the OOA envelope of each subject and condition. FOOOF allows for the examination of putative oscillations (peaks) in the frequency domain and characterizes these on their specific center frequencies, amplitude, and bandwidth by separating the periodic and aperiodic components of neural power spectra (Haller et al., 2018).

For statistical analyses of the periodic components of the OOA the attention modulation index (AMI) of both conditions was calculated using the following formula: (Attend Auditory – Attend Visual) / (Attend Auditory + Attend Visual) * 100. A two-tailed one sample t-test against 0 for each ear was calculated for the AMI pooled across the full range of the cochlear frequency response (1000-2000 Hz) and the range of extracted peaks from the left (3-10 Hz) and right ear (1-10 Hz). A nonparametric cluster-based permutation analysis over the whole brain was conducted to assess MEG-power effects in the cue-target period. The analysis was pooled across 1.7 s of the cue-target period and limited to a frequency range of 3-25 Hz. In a next step the AMI of the MEG-data was calculated and correlated with the OOA-AMI of the left and right ear. In order to assess statistical significance of the correlation, a nonparametric cluster-based permutation analysis over the whole brain was conducted. As for the assessment of MEG-power effects this analysis was pooled across 1.7 s of the cue-target period and limited to a frequency range of 3-25 Hz. The statistical analyses of the OOA and MEG data were conducted using the open-source FieldTrip toolbox for EEG/MEG data (Oostenveld et al., 2011), custom written MATLAB scripts (Version 9.1; The MathWorks), the R package “uniftest: Tests for Uniformity” (Melnik & Pusev, 2015), and custom written R scripts (Version 4.0.0; R Core Team).

## Results

### Behavioral Results

Performance was similar for both conditions and in general very high, underlining the compliance of the participants during the experiment. The average hit rates were *M* = 93.19 % (*SD* = 7.46 %) for the auditory task and *M* = 92.89 % (*SD* = 7.65 %) for the visual task. The hit rates of the two conditions did not differ significantly (*t*_(26)_ = 0.378, *p* = 0.709).

### OOA at Theta Rhythm Is Modulated by Intermodal Attention

Typical oscillatory activity of the brain is pronounced in a frequency band of 1-80 Hz, whereas otoacoustic activity is found at much higher frequencies (500-4000 Hz). As the aim of this experiment is to study the effects of cortical top-down modulations on OOA, we applied the Hilbert transform to extract the amplitude modulation for frequencies typical of ongoing cortical oscillations. To avoid a stronger influence of the lower sound frequencies and to create a representation of the cochlea’s frequency response, the otoacoustic signal was bandpass filtered between 1000 and 2000 Hz in 10 Hz steps with a window size of +/- 30 Hz. The PSDs of the 201 bandpass windows were then concatenated to create a representation of the amplitude modulation between 1000 and 2000 Hz of the cochlea’s frequency response.

In a first step we parameterized oscillatory modulations of OOA during the silent cue-target interval. We used FOOOF to differentiate between genuine oscillatory contributions from aperiodic 1/f changes. In all subjects a peak could be found at low (< 11 Hz) frequencies with a clustering around ∼5-6 Hz. For the Attend Auditory condition the average peak frequency was at 5.65 Hz (*SD* = 1.48) for the left and 5.88 Hz (*SD* = 2.33) for the right ear. For the Attend Visual condition the average peak frequency was at 5.58 Hz (*SD* = 1.57) for the left and at 5.85 Hz (*SD* = 1.83) for the right ear. Which modality was attended to had no statistically significant impact on the peak frequencies in both ears (left: *t*_(26)_ = 0.2068, *p* = 0.8378; right: *t*_(26)_ = 0.0681, *p* = 0.9462). **Figures 2A** and **2B** show subjects’ individual peak frequencies and **Figures 2C** and **2D** the slope for aperiodic components (“1/f noise”). Kolmogorov-Smirnov tests were performed to test for uniformity on the peak frequencies for every ear and condition. The percentage of peak frequencies for the left ear and Attend Auditory condition, *D*_*(26)*_ = 9.2347, *p* < 0.0001, and the percentage of peak frequencies for the left ear and Attend Visual, *D*_*(26)*_ = 9.2486, *p* < 0.0001, were both significantly different from uniformity, indicating that the peak frequencies were not uniformly distributed in both conditions. The same holds true for the right ear (Attend Auditory: *D*_*(26)*_ = 9.4619, *p* < 0.0001; Attend Visual: *D*_*(26)*_ = 9.3502, *p* < 0.0001).

**Figure 2.**
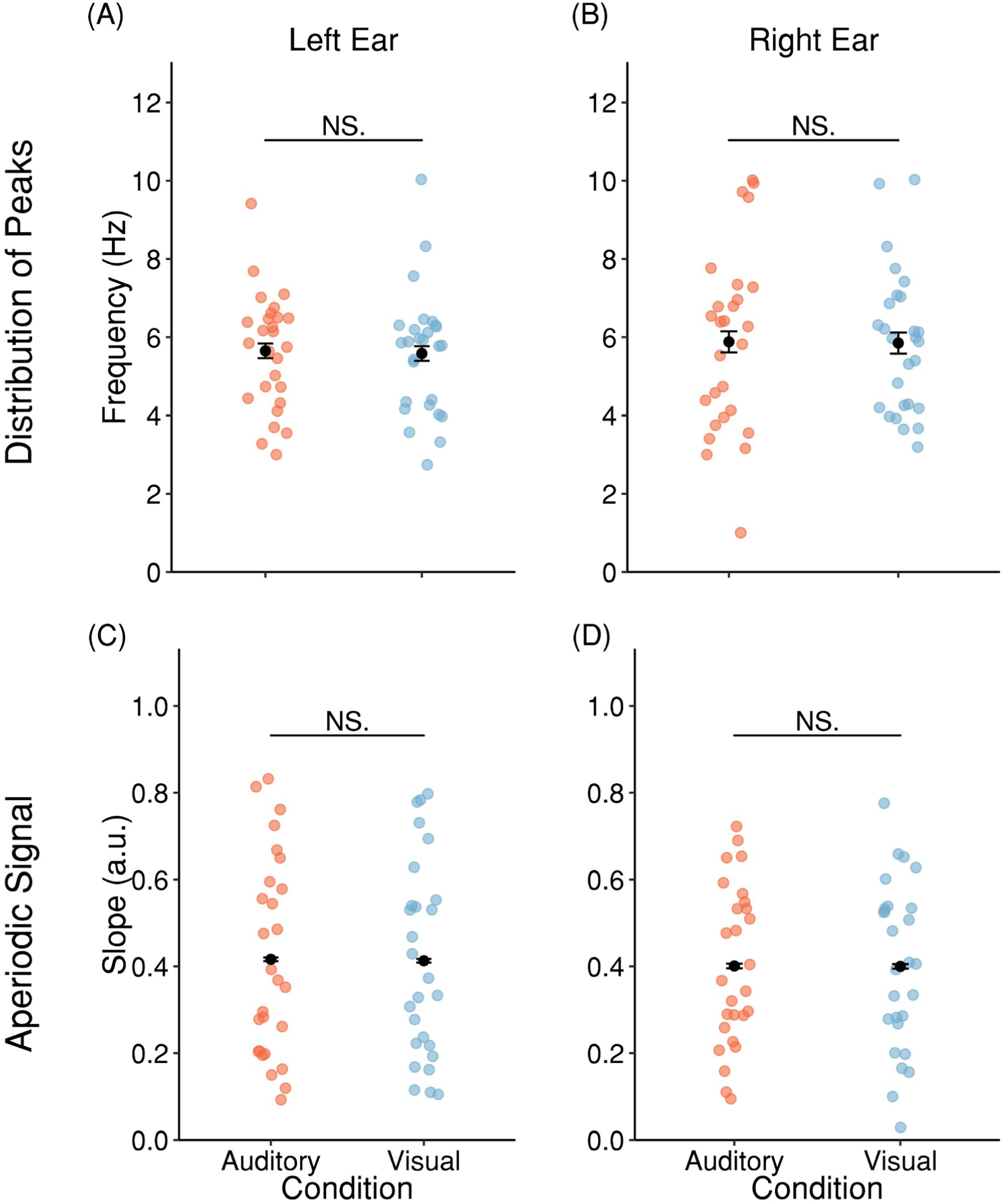
Peak analysis of OOA by FOOOF shows theta rhythmicity of cochlear activity. (A) Distribution of the peaks in the left ear for each subject and condition. (B) Distribution of the peaks in the right ear for each subject and condition. (C) Slope of the aperiodic signal in the left ear for each subject and condition. (D) Slope of the aperiodic signal in the right ear for each subject and condition. The black dots and error bars represent the mean and SEM (corrected for within-subject designs; see Cousineau & O’Brien, 2014).

While this analysis overall points to a theta-rhythmic modulation of cochlear activity in a silent cue-target period, the range (1-10.03 Hz) of these peaks suggests a rather high interindividual variability.

Next, we tested the hypothesis that cochlear activity is increased during periods of focused auditory compared to visual attention. Descriptively it appears from the grand average that the amplitude (**Figures 3A** and **3B**) differences of the AMI lie predominantly in the range of low frequencies, corresponding to the frequency range of dominant rhythmic cochlear activity (**Figures 2A** and **2B**). Given this overlap the AMI was pooled across the range of peak frequencies (left ear: 3-10 Hz; right ear: 1-10 Hz) for the cochlear response frequency range of 1000-2000 Hz for the left and right ear, respectively. In a next step one-tailed one sample t-tests against 0 were performed (see **Figure 3C**). The result for the left ear revealed that cochlear activity (*M* = 1.1002 %, *SE* = 0.3047 %) was significantly higher for the Attend Auditory condition (*t*_(26)_ = 2.4701, *p* = 0.0102). Similarly, the result for the right ear revealed significantly higher cochlear activity (*M* = 1.5343 %, *SE* = 0.3047 %) for the Attend Auditory condition (*t*_(26)_ = 2.3881, *p* = 0.0122). No interaural differences could be observed (*t*_(26)_ = -0.8225, *p* = 0.4183).

**Figure 3.**
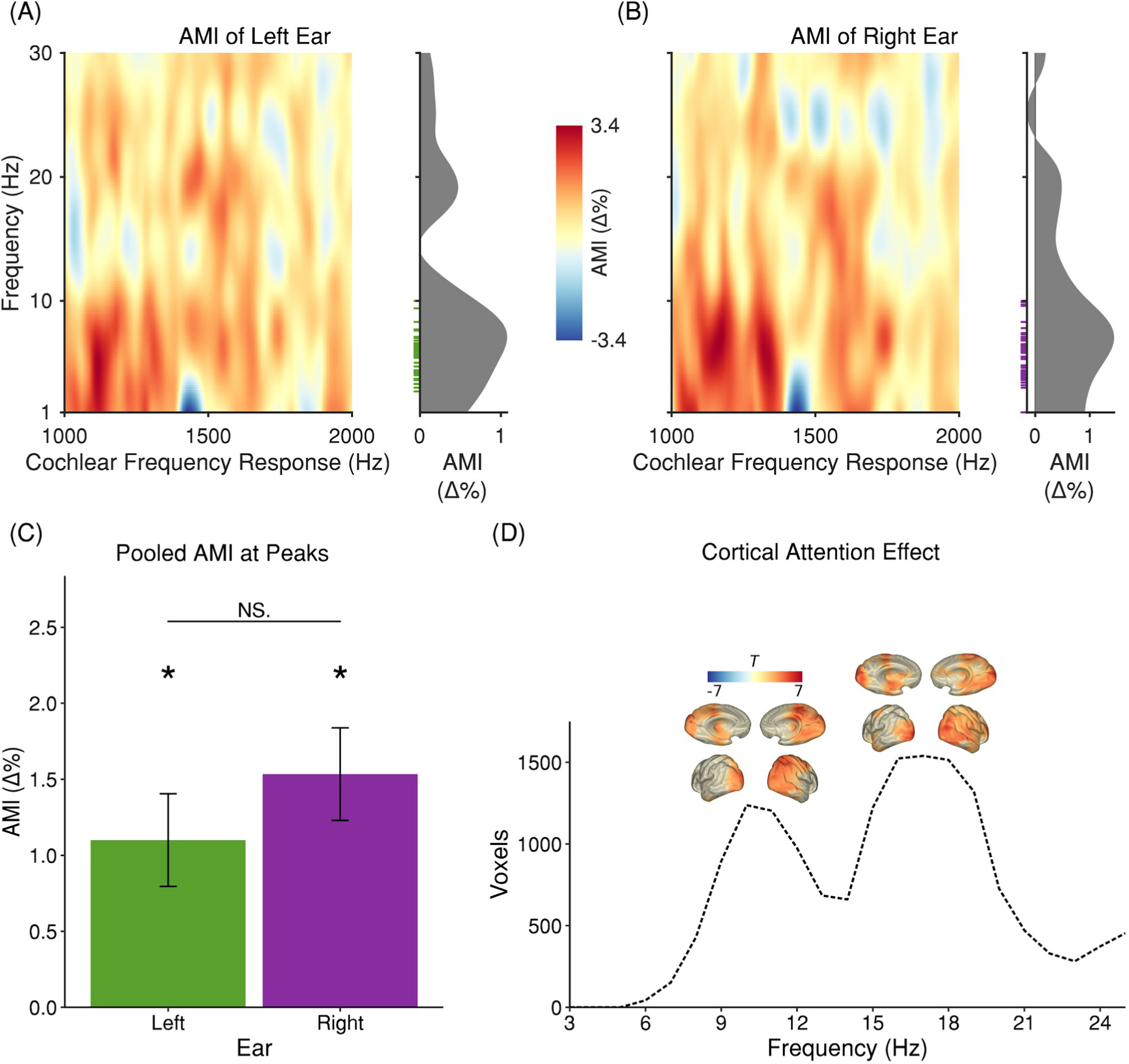
Power analysis of OOA shows enhanced low-frequency power for auditory attention. Power analysis of cortical activity reveals enhanced alpha- and beta-power for auditory attention. (A) & (B) AMI of the cochlear frequency response for the left and right ear. The x-axis represents otoacoustic activity at sound frequencies from 1000-2000 Hz. The y-axis represents the frequency range of the FFT. On the right of each subplot the OOA-AMI averaged over sound frequencies from 1000-2000 Hz is shown. The green and violet ticks illustrate the distribution of subjects’ peak frequencies from Figures 2A & 2B. (C) OOA-AMI averaged over sound frequencies from 1000-2000 Hz and the range of subjects’ peak frequencies (3-10 Hz for the left and 1-10 Hz for right ear). The OOA-AMI is significantly higher for the Attend Auditory condition in the left (*t*_(26)_ = 2.4701, *p* = 0.0204) and the right (*t*_(26)_ = 2.3881, *p* = 0.0245) ear. There was no difference between ears (*t*_(26)_ = -0.8225, *p* = 0.4183). (D) A nonparametric cluster-based permutation analysis indicated an effect of condition for brain power pooled across 0.25-1.95 s of the cue-target period (*p* = 0.004). This corresponded to a positive cluster in the observed data beginning around 4-6 Hz up to 24-25 Hz. The number of voxels in this cluster are shown as a function of frequency. The extent of the cluster is largest in the alpha- and beta-band. Moreover, for both bands it is located in posterior regions.

### Cortical Alpha and Theta Power Are Related to Cochlear Changes

In order to assess effects of intermodal attention on brain level, we performed a nonparametric cluster-based permutation analysis on source-projected MEG-power over frequencies of 3-25 Hz (see Materials and Methods section). The analysis was pooled across 1.7 s of the cue-target interval. An effect of condition (Attend Auditory > Attend Visual, *p* = 0.004) was observed that corresponded to a positive cluster in the observed data beginning around 4-6 Hz up to 24-25 Hz. As hypothesized, the extent of this cluster is largest in the alpha and beta range and located in posterior - mainly occipital and parietal-brain regions (see **Figure 3D**).

We expected inhibited sensory processing of the current task-irrelevant sensory modality – occipital regions for the visual and temporal regions for the auditory modality. According to dominant frameworks (Klimesch, 2012) this functional inhibition should manifest as increased power in the alpha-band. We found increased alpha power for the Attend Auditory condition over occipital regions. However, no increased alpha power for the Attend Visual condition in auditory regions could be found. This absence may be related to a reduced measurement sensitivity due to the significant loss of MEG sensors covering the temporal regions.

In order to assess whether attentional effects found at the cortical level were associated with the previously described cochlear effects, a correlation between the brain-AMI and the OOA-AMI of the left and right ear, respectively, was calculated. A nonparametric cluster-based permutation analysis indicated a significant correlation of brain-AMI and OOA-AMI of the right ear (*p* = 0.01) but not the left ear (*p* = 0.62). This corresponded to a negative cluster in the observed data incorporating the whole frequency range (3-25 Hz) of the analysis (see **Figure 4A**). The extent of the cluster peaks in the alpha-, theta- and beta-band. Dominant locations of the correlation effect are illustrated in **Figure 4A**. For the theta and alpha frequency range strong auditory cortical effects are seen in the left STG or medial portions of Heschl’s Gyrus, respectively. Interestingly the effects are strongest contralateral to the OAE probe. However effects were also observed outside of classical auditory cortical regions, such as in right (pre-motor) or left inferomedial temporal regions. To illustrate that effects are not driven by outlying participants of relevant effects in the theta- and alpha-band, **Figures 4B** and **4C** show correlations for voxels with the strongest effects. The negative correlations indicate that lower alpha- and theta-AMI is accompanied by higher OOA-AMI and vice versa. It is well known, that decreasing alpha-activity represents a mechanism for a release of inhibition (Jensen & Mazaheri, 2010; Klimesch, 2012). Thus, the negative correlation suggests that participants exhibiting a stronger release of inhibition (by lower alpha power) in left auditory brain regions during periods of auditory attention also exhibit elevated OOA-levels (by higher OOA power). This analysis illustrates that attentional modulations of rhythmic activity at the “lowest” (i.e. cochlear) level of the corticofugal system go along with modulations of oscillatory brain activity at the “highest” level.

**Figure 4.**
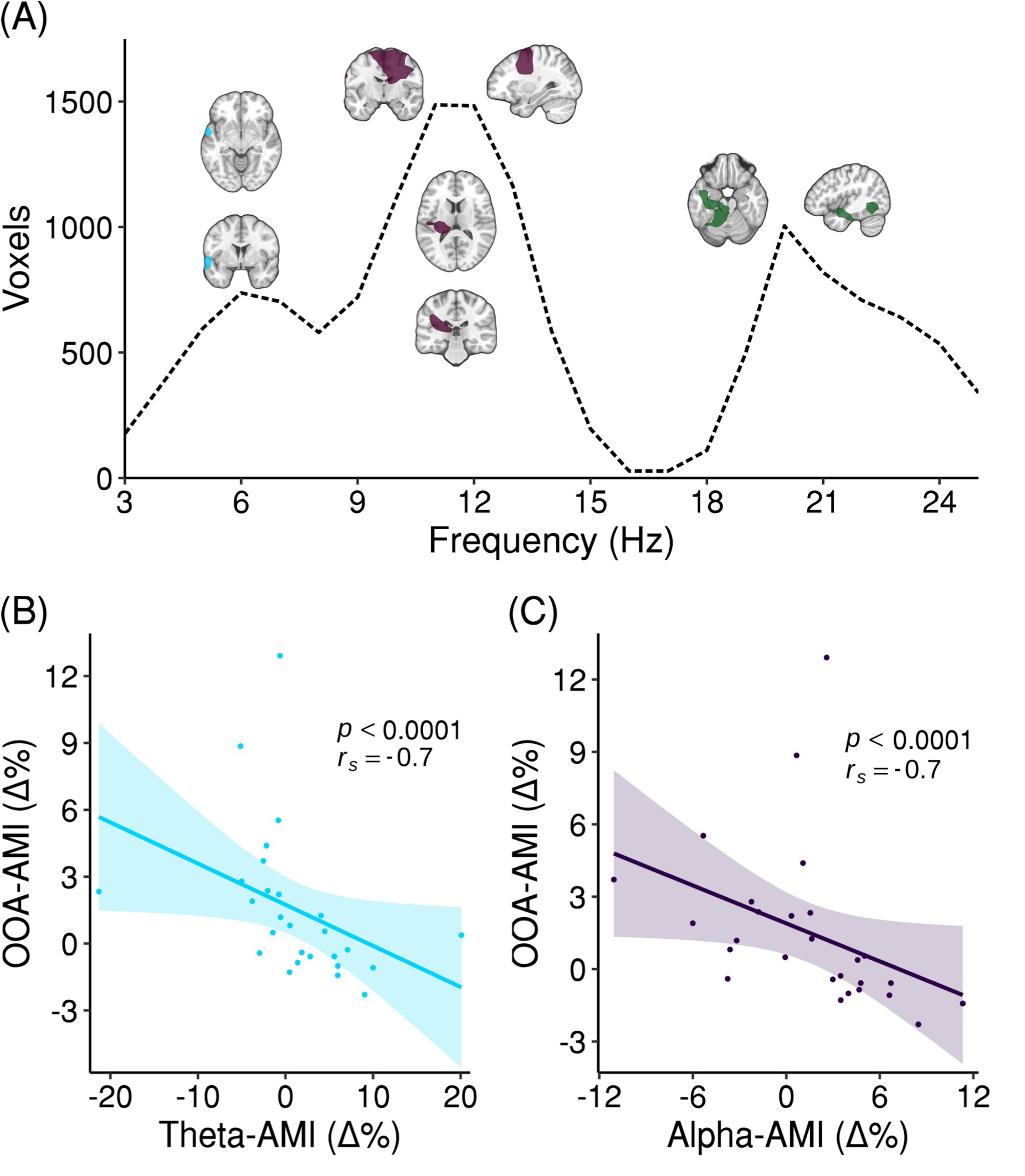
Correlation of cortical neural activity and OOA of the right ear. (A) A nonparametric cluster-based permutation analysis indicated a correlation of brain-AMI and OOA-AMI of the right ear pooled across 0.25-1.95 s of the cue-target period (*p* = 0.01). This corresponded to a negative cluster in the observed data incorporating the whole frequency range (3-25 Hz) of the analysis. The number of voxels in this cluster are shown as a function of frequency. The extent of the cluster peaks in the alpha-, theta- and beta-band. For the peak in the theta-band the cluster is located in the left STG. For the alpha-band it is located in medial portions of left Heschl’s Gyrus and right (pre-)motor areas. For the beta-band it is located in left inferior-medial temporal regions. Orthogonal views represent masked t-values (75 % threshold). (B) Correlation of brain-AMI at 6 Hz and OOA-AMI in the most significant voxel from (A). (C) Correlation of brain-AMI at 11-12 Hz and OOA-AMI in the most significant voxel from (A). The shaded error bars represent the SEM.

## Discussion

To what extent cochlear activity is sensitive to selective attention and how these changes are linked to cortical dynamics is a matter of ongoing debate. Given the uniqueness of the auditory system in having cortical descending projections from primary auditory cortex (via IC and SOC) to the cochlea, it is conceivable that a putative mechanism of alternating attentional states directly affecting cochlear processes could exist. To pursue our aims we adapted an previously introduced approach for investigating cochlear otoacoustic activity (Dragicevic et al., 2019) that allows us to draw first conclusions on how cortical attention processes are linked to cochlear otoacoustic activity. We demonstrate the presence of a theta-rhythmic pattern of otoacoustic activity during silent periods when attention was focused on either upcoming auditory or visual targets. Furthermore, we established a relationship between cochlear theta and cortical alpha modulations during the cue-target periods. Despite several open issues remaining, this study creates a connection between cochlear and cortical attentional modulations and helps close the gap between the remarkably segregated auditory attention research lines.

Our analysis of the OOA during the cue-target period indicated a genuine rhythmic modulation in the theta frequency range (∼6 Hz on average) that was not explicable by aperiodic (“1/f”) contributions to the spectrum. The peak frequency of the found rhythmic OOA pattern does not differ between visual and auditory attention, indicating that an endogenous cochlear rhythm at ∼6 Hz could exist. Depending on the generating mechanisms of the theta rhythmic cochlear activity, perceptual or attentional rhythmicities could either be genuine cortically driven effects (with cochlear effects being epiphenomenal) or they (and by extension cortical effects) could be an adaptation to cochlear physiological processes. However, the interindividual difference in peak frequencies was rather high, which hints at different mechanisms that putatively contribute to attention processes on the cochlea. This assumption is backed by the active sampling (Schroeder et al., 2010) literature, which points to the ubiquitousness of theta-like rhythms in various cognitive domains ranging from perception to action (Hasselmo & Stern, 2014; Poeppel, 2003; Spyropoulos et al., 2018; Tomassini et al., 2017). Extending such views, a recent “rhythmic theory of attention” framework states that attention is theta-rhythmically discontinuous over time (Fiebelkorn & Kastner, 2019; Fries et al., 2001; Landau & Fries, 2012; Wutz et al., 2018). While the latter framework has been developed mainly to better understand visuospatial attention, similar processes may also be relevant in the auditory system. For example (not in the focus of the current study), it is conceivable that interaural attention modulates the phase of the theta rhythm in both ears, facilitating signal transduction in the to-be-attended ear.

Beyond the illustration of a slow (theta) rhythmic modulation of OOA during silent cue-target intervals independent of the attention focus, we show that the magnitude of this process is clearly attentionally modulated. We found an enhancement during auditory selective attention, which might reflect an enhancement of cochlear sound amplification. In line with previous studies that found reduced levels of OAEs in subjects attending to a visual task, our results resemble an elevation of the to-be-attended acoustic stimulus during acoustic selective attention (Froehlich, Collet, Valatx, & Morgon, 1993; Meric & Collet, 1992; Puel, Rebillard, Bonfils, & Pujol, 1989; Wittekindt et al., 2014; see Smith et al. (2012) for an exception). Particularly, one study consistently reported similar amplitude modulations at low frequencies (< 7 Hz; Dragicevic et al., 2019). Yet, thus far, all studies on humans that have investigated effects of attention on the cochlea in cue-target periods utilized different types of evoked OAEs (EOAE) and distortion product OAEs (DPOAE). The measurement of EOAEs and DPOAEs relies on acoustic elicitor and probe stimuli, which are able to alter cochlear properties by themselves, making them rather unfavorable for assessing pure efferent effects (Guinan et al., 2003). It has to be noted that there are two studies that also investigated effects of attention (auditory & visual) and inattention on the cochlea by measuring physiological noise in a silent period subsequently of evoking nonlinear stimulus-frequency OAEs (Walsh et al., 2014a, 2014b). However, both studies differ from the current one as they analyzed cochlear activity after stimulation and did not compare auditory and visual attention effects. In our study, we utilized OOA that is measured in silent cue-target periods and therefore avoids any confounding efferent activity. Moreover, our approach allows us to stay as close as possible to previous literature in the cortical attention domain. In the current study we show power modulations of OOA in frequencies that in the cortical literature have been repeatedly reported to be related to various attentional task demands (Fiebelkorn et al., 2019; Fries et al., 2001; Klimesch, 2012; Wutz et al., 2018). Electrical stimulation of the auditory cortex in bats and chinchillas shows that cochlear responses can be modulated in a frequency specific manner (Dragicevic et al., 2015; León et al., 2012; Xiao & Suga, 2002). The current results imply that the modulation of cochlear low-frequency oscillatory power putatively is driven by top-down attentional processes (note that the frequency is unchanged). Given the well-established neuroanatomy of the auditory efferent system, corticofugal projections from the auditory cortex to the cochlear receptor, which are mediated by the IC and SOC, are the most probable neural substrates of this effect. The correlation effects of the present study, are compatible with this interpretation.

The current results of induced oscillatory activity in the MEG are in accordance with previous results and give an insight into the attentional demands of the task. Despite the unfavorable measurement conditions, we found elevated alpha- and beta-band activity in the pretarget period of Attend Auditory compared to Attend Visual trials in posterior regions but no modulations over auditory regions. Various studies on intermodal selective attention have postulated an active role of cortical alpha oscillations in modulating primary sensory areas (Bauer et al., 2012; Foxe et al., 1998; Frey et al., 2014; Fu et al., 2001; Wittekindt et al., 2014). In this context, alpha-band activity is proposed to reflect a suppression mechanism and especially seems to be relevant if distracting input has to be actively blocked. Two studies employing an audiovisual task have reported alpha power increases in posterior sensors when attention was directed to the auditory modality, power decreases when attention was directed to the visual modality, and no alpha-band modulations over auditory cortices (Foxe et al., 1998; Fu et al., 2001). In line with these findings, Wittekindt et al. (2014) observed a relative posterior alpha power increase when attention was focused on the upcoming auditory compared with the visual target. Our findings showing increased alpha power in primary visual cortex during auditory selective attention are in accordance with this view. In this way, alpha oscillations act to reduce processing of distracting input for the task-irrelevant visual modality.

Three previous studies have simultaneously recorded DPOAEs and EEG and were therefore able to investigate the relationship between cochlear and brain activity. Wittekindt et al. (2014) failed to show any correlations between those two. The authors explain this by the fact that their found effects depict different mechanisms of selective attention and thus do not depend on each other directly. In contrast, Dragicevic et al. (2019) reported significant correlations between the oscillatory DPOAE signal and cortical oscillations at low frequencies (< 10 Hz) mainly when attention was switched from the visual to the auditory modality. Finally, studying predictive processing using an intermodal predictability paradigm Riecke et al. (2020) found a relationship between DPOAE and brain effects. However, this relationship is limited to participants that benefited from predictions. Overall, as mentioned above, the elicitor stimuli which are required to evoke DPOAEs are prone to elicit MOC efferent activity that causes intrinsic cochlear changes by themselves. Hence, any inferences from correlations between oscillatory activity of the cochlea and the brain have to be treated with caution. The current study avoids these pitfalls by utilizing OOA in silent periods.

We found evidence for a putative relationship, namely, a negative correlation of cochlear low-frequency (1-10 Hz) power of the right ear and brain power, during periods of selective attention. This correlation was especially pronounced in the alpha-, theta-, and beta-band and was located in left auditory processing regions. It appears that subjects that exhibit a stronger cortical release of inhibition of auditory input (by reduced alpha-power) at the same time show stronger enhancement of the auditory target in the auditory periphery (by enhanced low-frequency OAA-power) and vice versa. Furthermore, the correlation in the theta-band is strongest at ∼6 Hz, the same frequency as the extracted periodic component of the OOA. Taking the relationships in the alpha- and theta-band together, they could point to a mechanism for a release of inhibition. Considering the architecture of the auditory efferent system it is likely that the outlined auditory cortical regions are a departure point for top-down modulations of cochlear activity in the current experiment. The observed cortico-cochlear correlations are compatible with the notion that these top-down modulations propagate through the efferent auditory pathway via crossed MOC fibres (Lopez-Poveda, 2018). Interindividual variability appears to exist to the extent that this top-down modulation is deployed next to the predominant inhibition of visual processing regions. In accordance with our findings (see also Dragicevic et al. (2019)) we suggest that top-down control of cochlear processing by cortical regions is mediated by slow oscillatory brain activity.

## Conclusion

The present study implies the existence of an putatively endogenous cochlear rhythm in the theta-band - a rhythm suggested to be linked to active sampling of the environment in different modalities (Fiebelkorn & Kastner, 2019; Landau & Fries, 2012; Schroeder et al., 2010). An outstanding question for future research is to understand the mechanistic relationship between cochlear theta rhythms and - especially auditory - cortical rhythms. Our results show that cochlear activity is modulated by intermodal top-down attention. In this regard, it provides evidence for the ongoing debate, whether the human auditory periphery is sensitive to top-down modulations (Beim et al., 2018; Lopez-Poveda, 2018). Future studies should investigate how these processes are manifested in individuals with reported hearing problems with or without audiometric deficits.

## Acknowledgments

We thank Mr. Manfred Seifter for his help with the measurements. The work of Moritz Herbert Albrecht Köhler is supported by a DOC fellowship of the Austrian Academy of Sciences.

